# Exploring phytoconstituents of *Juglans regia* to treat cervical cancer using integrative Bioinformatics

**DOI:** 10.1101/2023.06.15.545164

**Authors:** Riya Dua, Tulika Bhardwaj, Irshad Ahmad, Pallavi Somvanshi

## Abstract

Cervical cancer is the fourth most common malignancy among women, which also turns out to be the most common cause of death in women worldwide. Medicinal plants have traditionally been used to treat various diseases and disorders. The current study utilizes the molecular docking technique to investigate the anticancer potential of *Juglans regia* phytoconstituents against cervical cancer target proteins. This study includes the microarray dataset analysis of GSE63678 from the NCBI Gene Expression Omnibus database to identify differentially expressed genes. Furthermore, network biology approaches were employed to construct protein-protein interaction of differentially expressed genes. Next, the computation of topological parameters utilizing Cytohubba renders the top five hub genes (*IGF1, FGF2, ESR1, MYL9, and MYH11*). In addition, *Juglans regia* phytocompounds mined from the IMPPAT database were subjected to molecular docking analysis against identified hub genes. The application of molecular dynamics simulation validated the stability of prioritized docked complexes with minimum binding energy.

## Introduction

*Juglans regia* also known as English or Persian walnut, is a member of the *Juglandaceae* family exhibiting therapeutic potential against coronary heart disease, rheumatoid arthritis, cancer, and diabetes (Delaviz et al., 2017). It is densely cultivated in Asia, Central Europe, and the United States. It is considered a reputable source of nutrients and phytochemicals that greatly benefit human health, such as polyphenols, proteins, fibers, sterols, and essential fatty acids (Mohammadi et al., 2012). Several *in-vitro* studies validated the anticancer potential of *Juglans regia* with efficiency (Mohammadi et al., 2011). Natural phytocompounds, due to their lesser side effects and cost efficiency, are considered potential leads for drug discovery. Cervical cancer is the fourth most common cause of cancer and the fourth most common cause of death among women (Zhang et al., 2020). Cervical cancer is a female malignant tumor found in cervix cells located at the lower part of the uterus connecting to the vagina, caused due to abnormal growth of these cells (Song et al., 2013; Sundaram et al., 2020). The symptoms include vaginal bleeding, pelvic pains, and many more. Primary causes are HPV (human papillomavirus) infection, stress, smoking, and other STIs (Mattiuzzi and Lippi, 2020).

PPIs (protein-protein interactions) encompass many biological processes, including cell-to-cell communication, metabolic regulation, and development control (Sumanthy et al., 2014). PPIs have accelerated the modeling of functional pathways to illustrate the biological process’ molecular mechanism. PPIs can modify the kinetic properties of the enzymes, construct a new building site for small effector molecules, change the specificity of a protein for its substrate through interaction with different binding partners (Ajucarmelprecilla et al., 2022), or serve a regulatory role in either upstream a downstream level (Folador et al., 2016). A network represents a collection of points (proteins, genes.) joined together by lines and relationships between points. The network contains a set of nodes and edges. PPI network welcomes computation of topological properties of the underlying and computing modules, degree, and centrality of participating proteins. This results in the identification of ‘hub genes”. Hub genes are characterized as having a strong correlation in candidate modules where a high correlation or connectivity score is in the top 10%. For instance, if the module size is 1000, the top 100 genes with a high connection degree are referred to as hub genes (Dashti et al., 2020).

The current study aims to identify and analyze the therapeutic potential of phytocompounds from *Juglans regia* for treating cervical, endometrial, and vulvar cancer. A related microarray dataset (GSE63678) was mined from the NCBI Gene Expression Omnibus database (GEO). R package was used to identify differentially expressed genes (DEGs), assess gene ontology (GO), and evaluate pathway enrichment. The DEGs were integrated to construct a protein– protein interaction (PPI) network. Next, hub genes were recognized using the Cytoscape software and cytohubba (related plug-in). In accordance, phytocompounds of *Juglans regia* obtained from the IMPPAT (Indian Medicinal Plants, Phytochemistry And Therapeutics) database were subjected to ADMET, drug-likeliness, and physiochemical characterization. Molecular docking analysis of hub genes with potential screened phytocompounds enables the prioritization of efficient docked complexes. Further, MD simulation validates the stability of docked complexes, resulting in the exploration of potential lead molecules to serve as drug discovery precursors.

## Materials and Methodology

### 2.1. Dataset collection and pre-processing

The microarray dataset from the GEO database of GSE accession number GSE63678 related to cervical cancer, endometrial cancer, and vulvar cancer was mined, including samples into six categories (cervical cancer tissues/cells, normal cervical tissue/ cells, vulvar cancer tissues/cells, normal vulvar tissue/cells, endometrial cancer tissue/cells, normal endometrial tissue/ cells). Pre-processing initiates with data normalization, and the log was used to lower the values due to their wide range. Following sample clustering, Principal Component analysis was performed. Principal component analysis explains how genes are related or not related to one another via 2D graph representation. While clustering, a correlation matrix was generated with coefficients ranging between 0 to 1. It is an excellent way to predict gene functions because genes that share a biological process are frequently co-regulated utilizing heatmap (Miller et al., 2021). The LIMMA package in R enables gene expression differential analysis. The package supports two-color spotted array preprocessing. The DEGs are screened in LIMMA by building a linear model and estimating with the Bayes-T test (Abu-Jamous and Kelly, 2018). The p-values and differential expression statistics were calculated using empirical Bayes. To properly visualize the results of the DE analysis, a volcano plot was generated, highlighting only the top 20 genes based on their p-values and logFC values. R codes and related generated figures are available in **Supplementary File 1**.

### 2.2. Protein-protein interaction

Using the STRING database and a high confidence score of 0.700 as the cutoff condition, a PPI network was built to determine the degree of similarity between genes at default parameters. STRING’s network and enrichment facilities aid in thoroughly characterizing user gene lists and functional genomics datasets and creating and sharing highly customized and augmented protein-protein association networks (Szklarczyk et al., 2019). Each interaction depicts two non-identical proteins produced by a different protein-coding gene locus. Cytoscape is a freely available visualization platform that enables the computation of network topological parameters and hub gene identification (Shannon et al., 2003). The Molecular COmplex DEtection (MCODE) algorithm (version 1.5.1) is utilized (maximum depth = 100, node score = 0.2, and K-core = 220) for the computation of subnetworks and gene-enriched modules within the primary network (Bader et al., 2003).

### 2.3 Hub genes identification

Cytohubba, a Cytoscape plugin, was used to compute the MCC score for each node in the network (Chin et al., 2014). This study designated the genes with the highest MCC values as hub genes. As a result, these plugins aided in the identification of closely related genes. StringApp, another cytoscape app, was used to import the network directly from the STRING database online.

### 2.4 Identification of potential chemical leads

The Indian Medicinal Plants, Phytochemistry And Therapeutics (IMPPAT) database (Mohanraj et al., 2018), the most significant resource on phytochemicals of Indian herbs, was utilized to iterate the potential phytochemical extracts from the bark, flower, fruit, and leaf, seeds, root and stem of *Juglans regia*. A ligand library of 1004 phytochemicals from the IMPATT database and literature studies was generated to identify potential chemical leads having the potential to inhibit the prioritized hub genes. Deletion of phytocompounds was performed manually and resulted in 210 screened compounds. Lipinski’s rule of five (RO5) (Bhardwaj et al., 2019), a widely used drug-likeness measure, was used to screen the 210 phytochemical ligand library for potential druglike molecules. One thousand-four phytochemicals passed the R05 drug-likeness filter. The 3D structures of the druglike phytochemicals were then energy-minimized using the OpenBabel toolbox’s obminimize (Bhardwaj et al., 2022). Finally, OpenBabel was used to transform the ligands’ energy-minimized 3D structures.sdf to.pdb format.

### 2.5 ADME/T selection

The ADME/T (absorption, distribution, metabolism, excretion, and toxicity) qualities play a significant part in drug filtering when drug-likeness is determined by assessing current drug candidates’ physiochemical attributes and structural aspects. During drug designing, it is important to predict the situation and movement of a drug in the human body (Devi et al., 2023). Medicinal development costs can be decreased, and the process’ overall success rate can be increased by predicting the ADME/T characteristics of drug compounds before drug design. Drugs are transported into the circulatory system by absorption. The medicine is distributed when it crosses the cell membrane barrier and enters numerous tissues, organs, or bodily fluids (Alsulimani et al., 2022). The initial (parent) substance undergoes metabolism and is changed into new chemicals known as metabolites. Redox enzymes, often known as cytochrome P450 enzymes, process the majority of small-molecule drug metabolism in the liver (Kushwaha et al., 2021). Excretion is when a drug’s primary form and associated metabolites are expelled from the body. The drug’s toxicity also impacts the human body.Using admetSAR 2.0, screened compounds’ ADMET (Absorption, Distribution, Metabolism, Excretion, and Toxicity) profiles were created (Yang et al., 2019). This web-based application identifies the ADMET qualities of screened input compounds with structural and functional similarity. It contains manually curated data on known chemical compounds (Bhardwaj et al., 2019).

### 2.5. MOLECULAR DOCKING

The molecular docking of energy-minimized 3D ligand and target protein was carried out using AutoDock version 4.2 (Morris et al., 2009). The respective Python script prepared for ligands and protein structures from AutoDockTools was used to transform their 3D structures from.pdb format to. pdbqt format. By assessing the critical residues in the target proteins, such as the catalytic residues and substrate binding residues, which are crucial for the function and specificity of the considered proteases, the appropriate grid box specified by the search space center and search space size was manually determined for identified hub genes at the individual level (Khan et al., 2018; Kausar et al., 2022). With the active site in the middle of the grid box, the docking simulations were performed using the Lamarckian Genetic algorithm (LGA) (Ashraf et al., 2022; Sen et al., 2022). Each ligand’s binding energy conformation was calculated using the default settings of 2,500,000 energy assessments, 27,000 generations, 150 for population size, 0.02 for gene mutation rate, and 0.8 for crossover rate of 10 runs. Visualization was performed by using PyMOL (Rigsby et al., 2016).

### 2.6. Molecular Dynamic Simulation

MD simulations for the potential inhibitors of hub genes were performed to evaluate the stability of their protein-ligand complexes using GROMACS 5.1.5 and the GROMOS96 54a7 force field (Satyam et al., 2021; Kumari et al., 2021). The topology for the top inhibitors was generated using the Automated Topology Builder (ATB) version 3.0 (atb.uq.edu.au). After being positioned in the middle of a cubic periodic box, the protein-ligand combination was solvated by adding simple point charge (SPC) fluids. The system’s net charge was then balanced by adding Na+ and Cl-ions. Utilizing the steepest descent algorithm, energy was reduced. The system was then heated to 310 K during a 500 ps with 2 fs endless number of particle, volume, and temperature (NVT) simulation (Ahmed et al., 2021). The pressure was then increased to 1 bar during a 500 ps, 2 fs, constant number of particle, pressure, and temperature (NPT) simulation. Protein and ligand were both position-restricted in the simulations mentioned above. The position constraint was then released, and a production MD run was run for 100 ns with a 2 fs time step. The structural coordinates were saved every 2 ps during the MD simulation. The v-rescale temperature was maintained at 310 K. Using the Parrinello-Rahman pressure coupling approach, and the pressure was kept at 1 bar (Van Der Spoel et al.,2005). The temperature and pressure coupling’s time constants were maintained at 0.1 ps and 2 ps, respectively. The long-range electrostatic interactions were calculated using the particle mesh Ewald (PME) method with fourth-order cubic interpolation and 0.16 nm grid spacing.

In contrast, the short-range interactions were computed for the atom pairs within the cutoff of 1.4 nm. The LINCS approach restricted all bonds (Singh et al., 2019). Using GROMACS scripts, the trajectories derived from the MD simulations were then utilized to compute and assess the protein’s radius of gyration (Rg), root mean square deviation (RMSD), and root mean square fluctuation (RMSF) of the protein backbone Cα-atoms.

## Results

### 3.1. Identification of DEGs

The dataset from the GEO-NCBI database was analyzed using the R programming language to identify DEGs. LIMMA, DESeq, ggplot2, GEOquery, and heat map were among the R packages used for dataset analysis and visualization. The LIMMA package was utilized to analyze gene expression microarray datasets and a linear model to analyze bio-assays. Deseq2 performs internal normalization, calculating the geometric mean for each gene. The Geoquery package assisted in retrieving data directly from the GEO-NCBI database. The heatmap library aids in the visualization of gene expression and clustering. The table enlists p-values and logFC values for all differentially expressed genes. Gene IDs with logFC values greater than 0.5 were considered upregulated genes, resulting in 207 in this study (**Supplementary File 2)**.

### 3.2. PPI network construction

Following the discovery of the upregulated genes, the upregulated genes were subjected to a STRING database, resulting in a network displaying multiple gene connections. The network contains proteins with the highest degrees, essential for finding hub genes and interactions between different proteins. The StringApp plugin in Cytoscape was utilized for the network interaction visualization (Figure 1).

**Figure 1:**
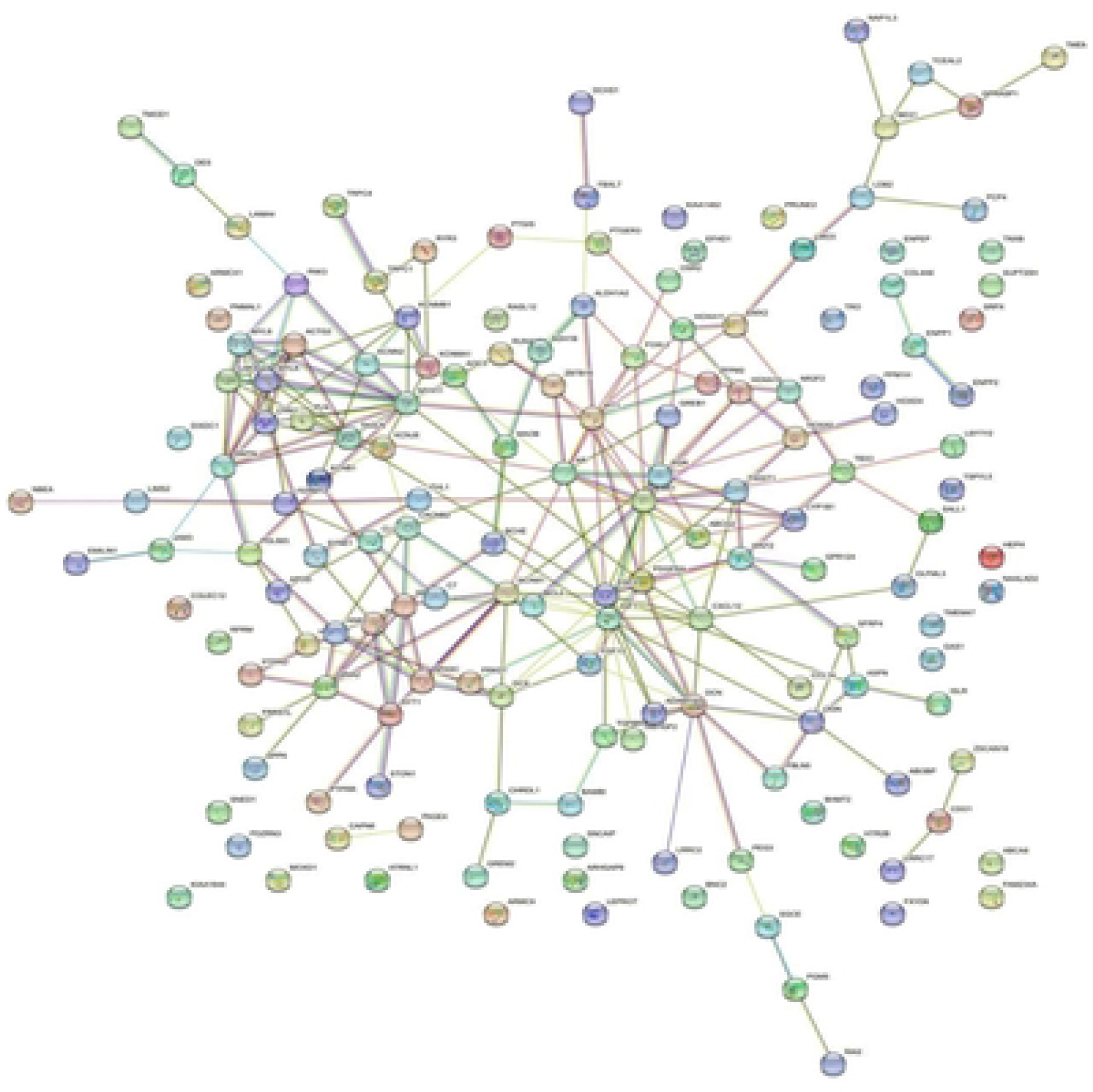
Visualization of Upregulated proteins networks

### 3.3. Identification of hub-genes

The generated PPI network had 207 nodes, 253 edges, 3.2 average node degree, 0.37 average local clustering coefficient, and 0.0903 PPI enrichment p-value. The created PPI network in this study contains much more acceptable interactions, according to the STRING database’s reference value (PPI enrichment p-value = 1.0e-16) (Figure 2). The interaction score of CytoHubba validated the top five hub genes. The CytoHubba outcomes of hub genes: Fibroblast Growth Factor 2*(FGF2)*, Insulin-like Growth Factor 1*(IGF1)*, Estrogen Receptor 1 *(ESR1)*, Myosin Light Chain 9*(MYL9)*, and Myosin Heavy Chain 11 *(MYH11)*.

**Figure 2:**
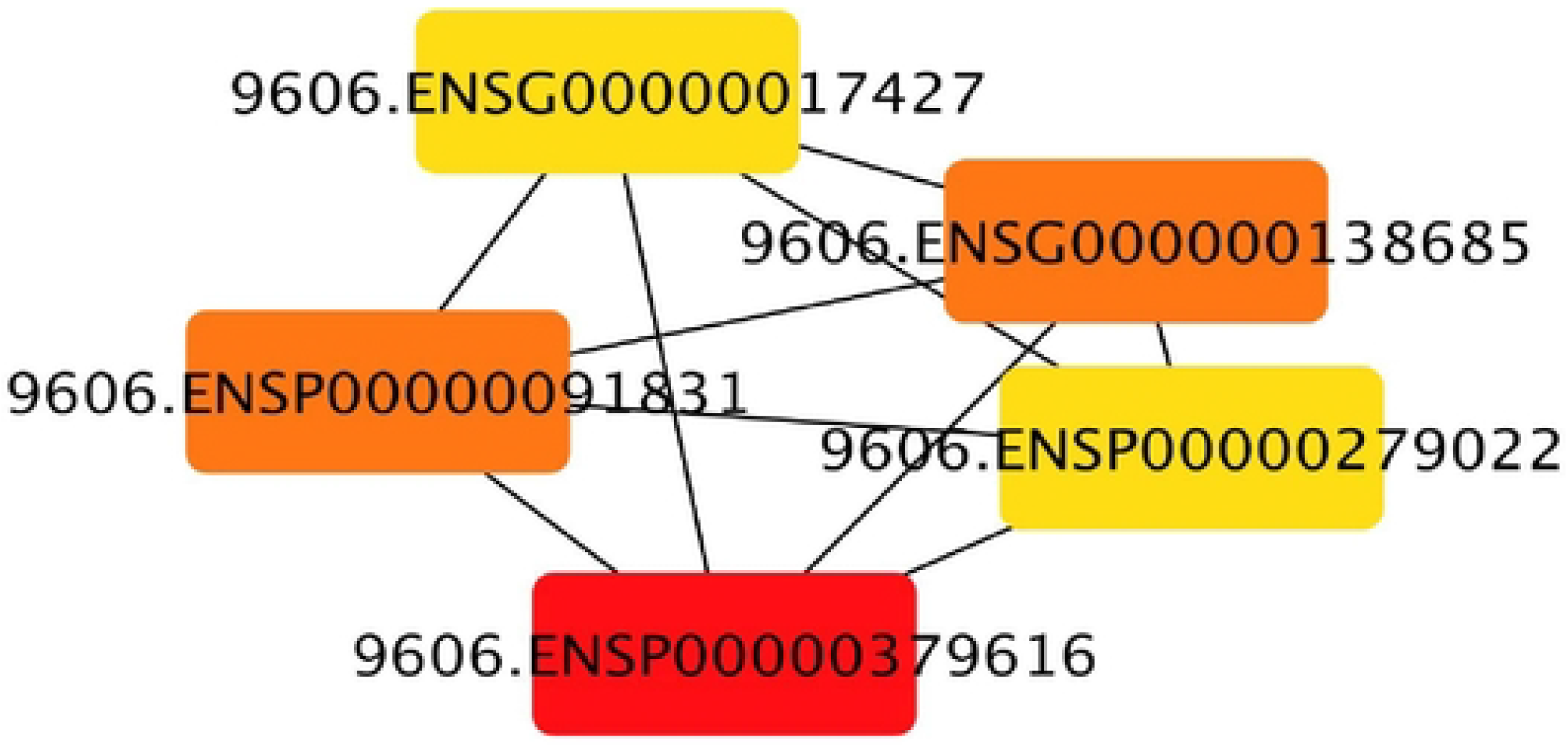
Visualization of lop five genes (ESRI. IGFI, FGF2, MYL9.MYH11) for cervical cancer dataset of 207 differentially expressed genes

### 3.4 Screening of potential phytochemicals

Lipinski’s rule of five (RO5) was used to compute the physicochemical parameters of the phytocompounds for five drug targets to assess their druglike characteristics. The molecular weight must be less than or equal to 500 g/mol, the number of hydrogen bond donors must be no more than 5, the number of hydrogen bond acceptors must be no more than 10, and the log p-value must be no more than five. One rule violation in a lead candidate is permissible. (**Supplementary File 3: Sheet 1**) displays the top hit phytochemicals and the reference compound’s anticipated druglike characteristics—all of the disclosed ligands displayed good druglike properties. Different pharmacokinetic features were predicted using admetSAR (The absorption, distribution, metabolism, elimination (ADME), and toxicity of the top medication candidate molecules can be predicted using pharmacokinetic variables. (**Supplementary File 3: Sheet 2**) displays both targets’ ADMET traits of the generated phytochemicals. Due to their poor pharmacokinetic qualities and toxicity, many medications must incorporate this method in their drug development. Early drug discovery relies on high-performance and quick ADMET profiling assays to identify active lead compounds (Bhardwaj et al., 2018). According to the

ADMET profiling, all of the candidate compounds had no adverse effects upon absorption. The associated ADMET and physiochemical properties (**Supplementary File 3: Sheet 3)** of potential compounds for different models, such as P-glycoprotein substrates, BBB penetration, and gastrointestinal absorption, showed positive results strongly support the compounds’ suitability as drug candidates.

### 3.5 Molecular Docking

The intermolecular interactions among proteins and ligands were analyzed for computing binding energies of protein-ligand complexes using AutoDock v 4.2 40. The structure-based virtual screening prioritized FGF2-myrcene, IGF1-Juglone, and FGF2-Juglanin as prioritized docked complexes. PubChem database was mined for the retrieval of chemical structures of the phytocompounds. The binding interactions of drug targets with their respective prioritized ligands are depicted in Figure 3 and Table 1. Contacting residues in each docked complex is also indicated.

**Figure 3:**
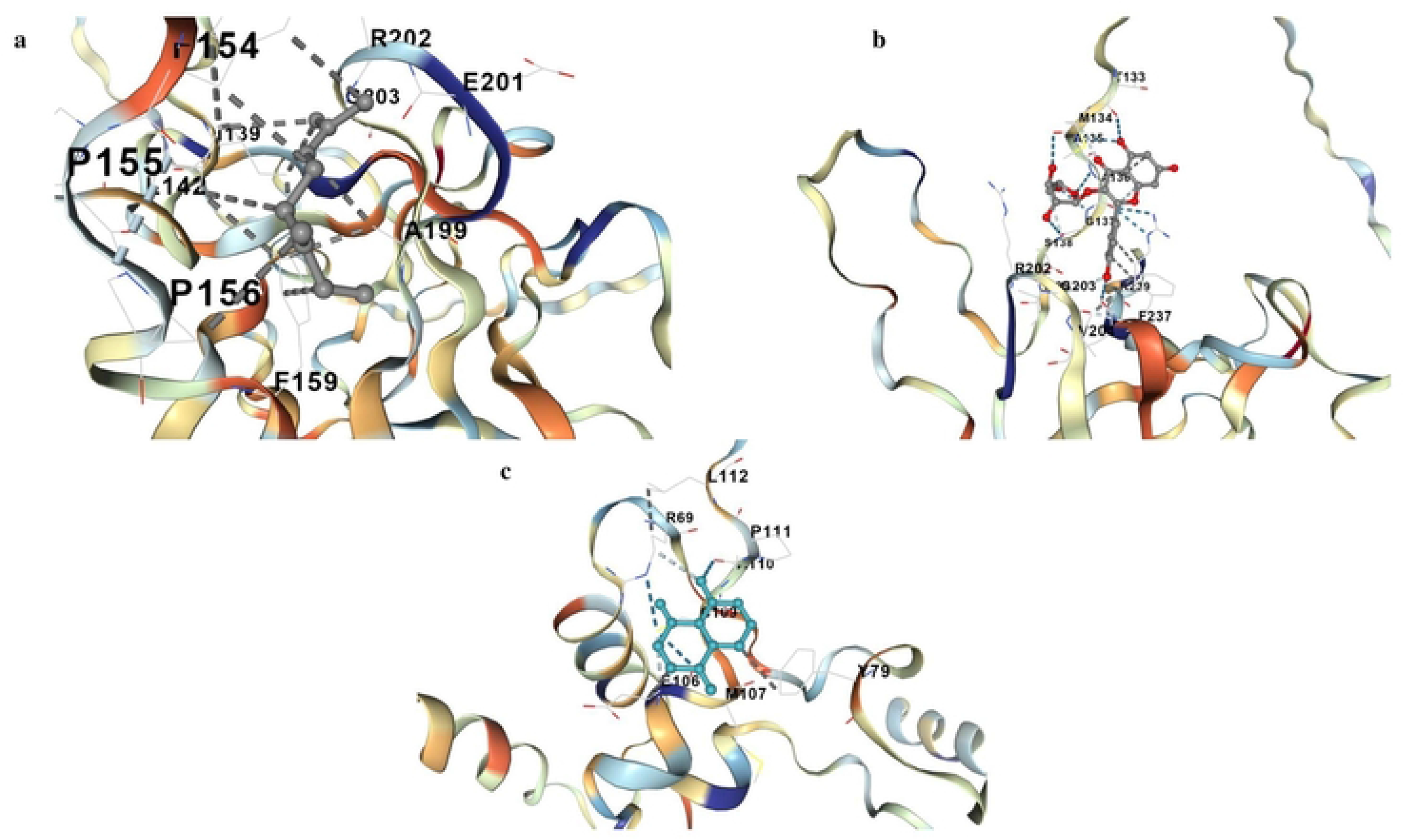
Visualization of docked complex of (a)FGF2-Myrcene. (b) FGF2-Juglanin and (c) IGFl-Juglone

## 3.5. MD simulation

To investigate the stability of the identified inhibitors’ protein-ligand complexes, MD simulations of 100 ns were performed for the top three protein-ligand docked complexes (FGF2-myrcene, IGF1-Juglone, and FGF2-Juglanin) by analyzing radius of gyration (Rg), root mean square deviation (RMSD), and root mean square fluctuations (RMSF) of the Cα atoms. Throughout the MD simulation, the Rg value of protein-ligand docked complexes remains largely stable (Figure 4a). The average Rg value of (FGF2-myrcene), IGF1-Juglone, and FGF2-Juglanin are 2.008± 0.003 nm, 1.962± 0.024 nm, and 2.313± 0.043 nm, respectively. Furthermore, after 20 ns, the RMSD value of the Cα atoms of docked complexes becomes stable (Figure 4b).*FGF2*-myrcene, *IGF1*-Juglone, and *FGF2*-Juglanin have an average RMSD value of 0.413± 0.012 nm, 0.487± 0.011 nm, and 0.618± 0.018 nm, respectively, over the 20 ns to 100 ns time interval. Finally, Figure 4c depicts the RMSF value per residue in FGF2-myrcene, IGF1-Juglone and FGF2-Juglanin complex.

**Figure 4:**
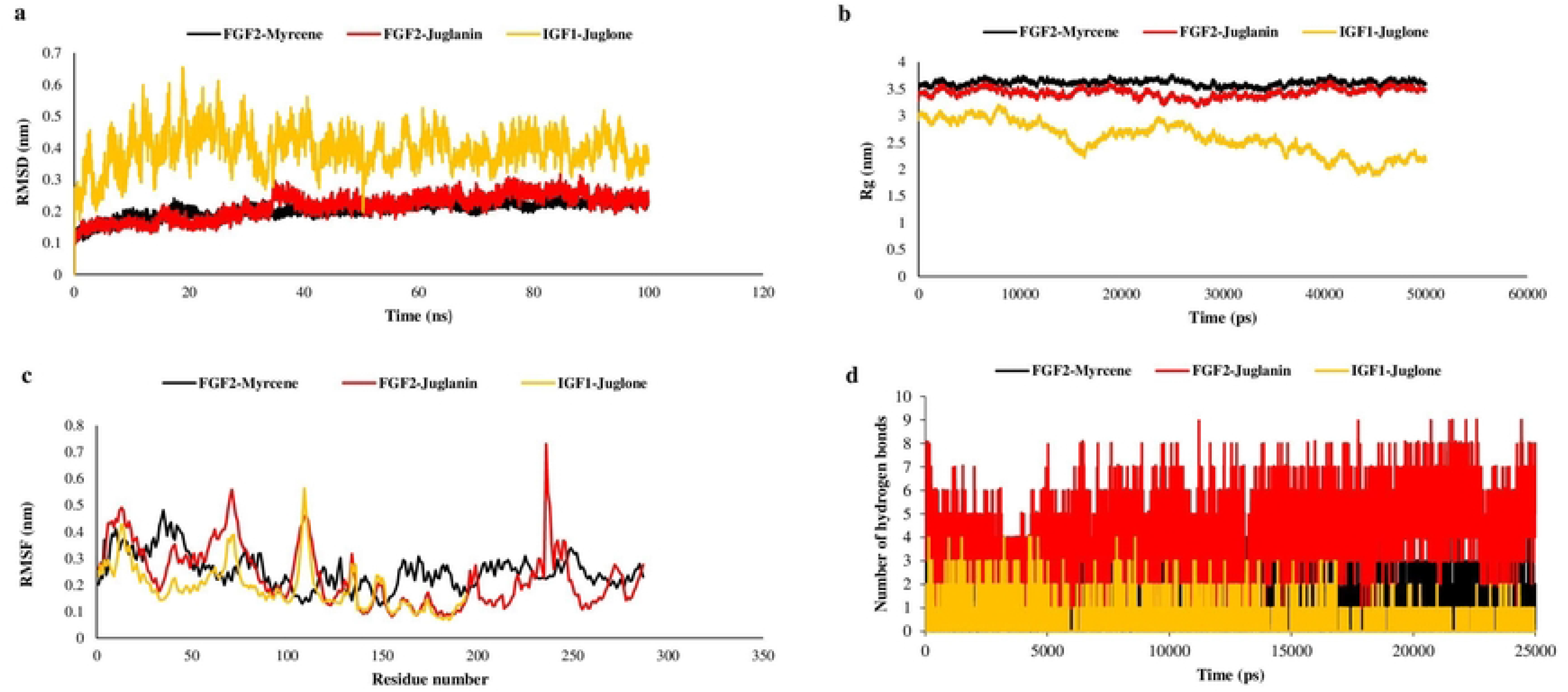
(a)RMSD, (b) Radius of Gyration, (c) RMSF and (d) Hydrogen bond analysis of FGF2-Myrcene, FGF2-Juglanin and 1GF1 -Juglone

## 4. Discussion

In the entire world, cervical cancer affects women more frequently than any other cancer. Primary prevention and screening are the best strategies for reducing the cost of care and mortality linked to cervical cancer. Therefore, phytoconstituents from herbs were considered for the drug discovery targeting cervical, vulvar, and endometrial cancers. These compounds possess enormous structural and chemical diversity with fewer side effects.

Although numerous studies have used microarray-based technology to find molecular markers in cervical, endometrial, and vulvar cancers, there have been differing results reports because of patient selection, tissue source, and study designs. Therefore, the present study attempts to identify the prioritized gene signatures underlying cervical, endometrial, and vulvar cancers by a comprehensive meta-analysis of the microarray dataset GSE63678. It identifies 412 differentially expressed genes (207 upregulated and 205 downregulated genes). By uncovering upregulated genes, this study highlights the potential diagnostic and prognostic biomarkers in female reproductive-related cancers and assists in understanding the molecular mechanism of their development and progression. Protein-protein interactions (PPIs) offer various biological processes, cell-to-cell interactions, and metabolic and developmental control (Rao et al., 2014). The elucidation of protein interaction networks also contributes significantly to analyzing network robustness and stability (Lu et al., 2020). Topological parameters computation of the network enables the identification of hub genes. It offers the advantage of extracting information from large volumes of high-dimensional data, which assists in identifying novel players involved in multiple proteomic interactions in cervical, endometrial, and vulvar cancer patients. This renders the top five differentially expressed hub genes (Fibroblast Growth Factor 2 *(FGF2)*, Insulin-like Growth Factor 1*(IGF1)*, Estrogen Receptor 1 *(ESR1)*, Myosin Light Chain 9*(MYL9)*, and Myosin Heavy Chain 11 *(MYH11))*. Fibroblast growth factor 2 protein encoded by the FGF2 gene involves various biological processes like tissue repair, embryonic development, tumor growth, and cell survival activities (Lou et al., 2014). It is experimentally validated as a potential target for treating gastric ulcers and myelofibrosis (Nunes et al., 2016). Single-chain polypeptides with a high degree of sequence homology to pro-insulin are insulin-like growth factors 1 and 2 (Yuan et al., 2018). It plays a crucial role in the growth and development of many tissues and regulation of overall growth and metabolism. It is an attractive target for treating cervical cancer, diabetes, and inflammation (Arcaro et al., 2013). Estrogen Receptor 1 *(ESR1)* is an attractive drug target for treating breast cancer, myocardial infarction, and migraine (Jeong et al., 2021; Brett et al., 2021). Myosin light chain 9 (*MYL9*) and myosin heavy chain (MYH11) plays a vital role in immune infiltration, tumor invasion, and metastasis, therefore, serve as a potential target for cancer treatment (Kandler et al., 2020).

Further, the chemical compounds of *Juglans regia*, mined from the IMPPAT database, were subjected to ADMET analysis. Screened non-toxic compounds were docked against identified plausible hub genes to identify selective inhibitors with optimal binding energy. This results in identifying phytochemicals from *Juglans regia*, which will serve as potential leads for drug discovery. Molecular docking analysis prioritizes myrcene, juglone, and juglans as attractive potential chemical lead capable of inhibiting Insulin growth factor 1 and Fibroblast Growth Factor 2. Myrcene, a yellow-colored oily liquid with a flash point of less than 200 °F is insoluble in water (Surendran et al., 2021). It is an octa-1,6-diene monoterpene with methyl and methylene substituents at positions 3 and 7, respectively, with anti-inflammatory properties (Polec et al., 2021), anti-aging and analgesic properties (Behr et al., 2009). Myrcene, extracted from the fruit of *Juglans regia*, is distributed in adipose tissues, liver, kidney, and brain (Hanus et al., 2020) with bioavailability of 30min in human plasma (Rufino et al., 2015). Several studies have documented the antibacterial effects of myrcene against gram-positive bacteria (Mahizan et al., 2019), for example, *Staphylococcus aureus* (Inoue et al., 2004), *Escherichia coli* (Galgano et al., 2022), *Salmonella enterica* (Ebani et al., 2019), etc. Myrcene can change the lipid monolayers’ fluidity, stability, and morphology (Polec et al., 2020). Myrcene exhibits antitumor activity against lung cancer cells by inducing oxidative stress and apoptosis (Bai et al., 2020). In the FGF2-myrcene complex, a binding energy of -9.17 kcal/mol was computed. ILE139, PHE154, ALA199, and GLY203 interact with methylene substituents at position 7. PRO155, ARG202, and GLU201 form hydrogen bonds with C3 and C5 positions.

Juglone (5-hydroxy-1,4-naphtoquinone), an oxygen derivative of naphthalene, is extracted from the leaf of *Juglans regia*. It is transformed into a toxic compound when exposed to soil or air. Therefore, an appropriate dosage (13.1–1556.0 mg/100 g) is required for drug design. Juglone exhibited anticancer effects on the different breast (Zhang et al., 2015), lung (Fiorito et al., 2016), prostate (Kanaoka et al., 2015), cervix (Zhang et al., 2012), and blood cancer models (Zhang et al., 2012). Juglone induced early DNA single-strand damage on human fibroblasts, translating into apoptosis and necrosis (Paulsen et al., 2005). Juglone inhibited metastasis development in the same cell line and decreased spheroid invasiveness (C6) (Meskelevicius et al., 2016). Juglone blocks several molecular pathways involved in cancer development, such as the PIK3/Akt cascade mechanism (Chae et al., 2012). IGF2-Juglone docked complex exhibits a binding energy of -9.02 kcal/mol. CYS109, LEU112, and GLU106 residues of IGF2 interact with the hydroxyl group at the C5 position. TYR79 and PRO111 residues are bonded with the C4 and C5 positions of juglone via hydrogen bond. The presence of van der Waals interactions with TYR79 and ALA110 with the C4 benzene ring group are essential structural requirements for anti-toxicity, also found during the analysis.

Juglanin is a polyphenol extracted from the pericarp of *Juglans regia*, exhibiting anticancer and anti-inflammatory properties. Apoptosis and autophagy were triggered simultaneously and exerted a synergistic activity when treated with Juglanin (Chen et al., 2017). Juglanin enhanced the effect of doxorubicin, one of the most commonly used antitumor drugs (dox). It significantly increased the cytotoxic effect of doxo in normal and doxo-resistant A549 cells and normal and cisplatin-resistant H69 cells (Wen et al., 2017). MAPKs, the proteins that regulate cellular proliferation, differentiation, and apoptosis, are considered an attractive target of Juglanin (Liu et al., 2005). Docking analysis revealed that MET134, ALA136, SER138, GLY203, and PHE237 play a significant part in the binding of the C-7 position of the hydroxyl group in the benzene ring. Primary H-bonding interaction with the benzene ring (C-5th position of –OH group) could associate with the (O-atom) key chain of GLU238, MET134.

Molecular dynamics simulations were used to examine the stability of protein-ligand complexes further. The protein target’s backbone structural framework generated root mean square deviation (RMSD) graphs for time at 100 ns. An average RMSD of 0.511 ± 0.21 nm was computed. RMSD values gained until 4 ns in the case of FGF2-Juglanin Fig. 4 (b) and IGF2-Juglone Fig. 4 (c) related complexes, and conjunction was detected from 100 ns, but slight variations remained throughout. Figures 4(a, b, and c) show overlaid graphical demonstrations of time-dependent RMSD of protein-ligand complexes (a). Most binding sites are thought to be shaped like alpha-helix, according to the Dictionary of Secondary Structure of Proteins (DSSP), and active sites are thought to be located in the coil region. To understand the flexibility of each residue, protein-ligand docked complexes were studied using residue-based root mean square fluctuations (RMSF). Due to the lack of structural data for the target protein, RMSF values for the initial 15 residues of each docked complex vary considerably (0.42-2.87 nm for each case). Less turbulence at the binding and active site implies that the binding cavity is rigid and intact. The docked complexes’ solidity and structural changes are calculated by gmx gyrate’s Rg values. It also determines the atomic mass corresponding to the mass centers of the complexes. With no variations after 50000 ps, the average Rg values of *FGF2*-Myrcene, *IGF2*-Juglone and *FGF2*-Juglanin were (2.36-2.66 nm), (2.29-2.89 nm), and (2.44-2.56 nm), respectively. Furthermore, the firmness of the prioritized protein-ligand complexes is confirmed by the correlation between Rg values and RMSD values of backbone C atoms.

## 5. Conclusion

Medicinal plants are potential precursors for drug discovery rather than synthetic compounds due to the chances of lesser side effects. Juglans regia is a traditional medicinal herb exhibiting antidiabetic, anticancer, and anti-inflammatory properties. Therefore, this study aims to identify potential phytocompounds with plausible inhibitory potential against cervical cancer-related drug targets. We identified specific critical genes utilizing protein-protein network analysis and topological parameter computations. *This* renders potential five genes subjected to molecular docking analysis and MD simulation studies with prioritized phytocompounds. Such compounds will be utilized for drug discovery against cervical cancer. Therefore, it can improve the survival rate and reduce the death rate, which concentrates on suppressing or controlling this gene’s function.

## Author Contributions

RD, TB acquired data, analysis, interpretation of data, and manuscript drafting. TB, IA, and PS study concepts and design, data acquisition, analysis and interpretation of data, drafting of the manuscript, and supervision. PS and IA did data interpretation; manuscript drafting. All the authors reviewed and approved the manuscript.

## Funding

PS is grateful to DBT grant no BT/PR40251/BTIS/137/11/2021 for providing the necessary infrastructure. The authors would like to thank Jawaharlal Nehru University and the Department of Biotechnology for the facility and fellowship for the requisite support. IA is thankful to the Deanship of Scientific Research at King Khalid University, Kingdom of Saudi Arabia (RCAMS/KKU/0018-22) for their support.

## Acknowledgments

The authors thank the Jawaharlal Nehru University facility and funding agency for supporting us.

## Conflicts of Interest

The authors declare no conflicts of interest.

## Notes

### Competing Interest Statement

The authors have declared no competing interest.

